# Optimizing Antimicrobial Susceptibility Testing: Cost and Environmental Benefits of MIC Volume Reduction

**DOI:** 10.1101/2025.04.29.651209

**Authors:** Jonathan Clarhaut, Jeremy Moreau, Tom Collet, Emma Babiard, Vincent Aranzana-Climent, Sandrine Marchand, Kevin Brunet, Julien M. Buyck

## Abstract

The determination of Minimum Inhibitory Concentrations (MICs) is essential for evaluating antimicrobial efficacy, guiding both clinical treatment decisions and drug development. The standard broth microdilution method is widely used but requires significant reagent volumes, which can be limiting when working with novel or expensive antimicrobials. This study assesses the feasibility of reducing assay volumes without compromising MIC accuracy. We compared the MIC values obtained in standard 96-well plates (200 µL) to those in 384-well plates with reduced volumes (30–50 µL) for a range of Gram-negative and Gram-positive bacteria, as well as yeast species. Our results demonstrate that except for micafungin, MIC values obtained with reduced volumes remained within the acceptable variability ranges defined by EUCAST and CLSI. Evaporation, a potential source of bias in smaller volumes, was mitigated by conducting experiments in a water-saturated atmosphere. Furthermore, reduced assay volumes significantly lowered material costs and antimicrobial consumption, particularly for expensive drugs such as cefiderocol. This miniaturization approach offers a cost-effective, high-throughput alternative for antimicrobial susceptibility testing while maintaining accuracy and reproducibility.

## Introduction

The determination of Minimal Inhibitory Concentration (MIC), which refers to the lowest concentration of an antimicrobial agent required to inhibit visible growth of a microorganism, plays a crucial role in the evaluation of antimicrobial efficacy, guiding both clinical treatment decisions and the development of novel therapeutics. Among the various methods for determining MIC, the broth microdilution technique has gained prominence due to its accuracy, reproducibility, and adaptability in a laboratory setting, and it is now generally considered as the gold standard method by institutions such as the European Committee on Antimicrobial Susceptibility Testing (EUCAST)(1) or the Clinical and Laboratory Standards Institute (CLSI) (2) in the USA.

Generally performed in 96-well microtiter plates, the microdilution method enables the testing of multiple antibiotic or antifungal concentrations simultaneously, thereby providing a comprehensive antimicrobial profile for both established and novel compounds. This method is generally automated in clinical laboratories but not typically in microbiology research laboratories. Based on the recommendations of the CLSI or the EUCAST, this method is applied in a final volume of 100 to 200 µL in accordance with ISO 20776-1 (3).

Although wholly standardized in terms of the medium used, microorganism inoculum, incubation time and temperature, in the context of testing new molecules or molecules synthesized in small amounts, or by using expensive broths like Iron-Depleted broth used to test the newly marketed cephalosporin siderophore cefiderocol, the reproducibility and reliability of this method in smaller volumes needs to be validated. By employing smaller volumes, microdilution not only conserves resources (by reducing plastic consumption) but also enhances experimental efficiency, reducing costs and increasing throughput. This makes it especially beneficial for laboratories working with large panels of antimicrobials or conducting high-throughput screening for drug development.

In this article, we conducted a comparative study on MIC results when testing volumes were reduced. To do so, the values of MIC done in 200, 100, 50 and 30 µL final volume in two different microtiter plates (96 and 384 wells) were compared with the reference strains for five different bacterial species (*Pseudomonas aeruginosa, Staphylococcus aureus, Escherichia coli, Klebsiella pneumoniae* and *Acinetobacter baumannii*) as well as two yeast species (*Candida krusei* and *Candida parapsilosis*) and with different antimicrobials (ciprofloxacin, gentamicin, linezolid, meropenem, amikacin, ceftazidime, aztreonam, amphotericin B, fluconazole, micafungin) representative of the main family used in clinics.

## Materials and Methods

### Strains used

The following reference strains were used for the study: *Pseudomonas aeruginosa* ATCC 27853, *Staphylococcus aureus* ATCC 29213, *Escherichia coli* ATCC 25922, *Klebsiella pneumoniae* ATCC 43816, *Acinetobacter baumannii* ATCC 19606, *Candida krusei* ATCC 6258 and *Candida parapsilosis* ATCC 22019.

### Antibiotics and reagents

Amikacin and gentamicin were provided from Acros (Thermo Fisher scientific, Illkirck-Graffenstaden, France) and Carl Roth (Lauterbourg, France) respectively. Amphotericin B, Aztreonam, ceftazidime, ciprofloxacin, fluconazole, linezolid, meropenem and micafungin were purchased from Sigma-Aldrich (Saint-Quentin Fallavier, France). Cefiderocol was supplied as Fetcroja by Shionogi.

Cation-adjusted Mueller Hinton Broth 2 (Ca-MHB) and RPMI 1640 (with L-glutamine, pH indicator, no bicarbonate supplemented with 2% (w/v) glucose and buffered to pH 7 with 0.165 mol/L MOPS) were purchased from Sigma-Aldrich.

Iron-Depleted MHB (ID-MHB) was homemade from Ca-MHB according to the European Committee on Antimicrobial Susceptibility Testing (EUCAST) recommendations (4). The concentration of Ferric ion was then checked by Inductively Coupled Plasma Mass Spectrometry (ICP-MS) and reached 8.29 µg/L for this batch.

### Evaluation of evaporation

To ensure repeatability, all experiments were conducted using the Assist Plus robot and automated pipettes (Integra Biosciences SAS, Cergy Pontoise, France). Two types of plates were used: 96-well plates for tests with 100 and 200 µL final volume and 384-well plates for tests with 30, 50 and 100 µL. To evaluate the impact of evaporation, plates with different final volume of Ca-MHB were incubated for 24 h in air atmosphere (plates were deposited in incubator at 37°C without agitation) or in water-saturated atmosphere (plates were placed in a box with water reservoir and in the incubator at 37°C without agitation) without addition of bacteria. The content of each well was weighed to measure the remaining volume after 24 hours of incubation at 37 ± 2°C, either in an air or a water-saturated atmosphere.

### MIC in microdilution

Minimum inhibitory concentrations (MICs) were determined following guidelines for standard conditions, with adjustments made as follows:

#### For the influence of the atmosphere

MIC was evaluated in 200 µL final volume in both atmospheres for the five bacterial species against different antibiotics as detailed in **Table 1**.

**Table 1.** Conditions tested in MIC to evaluate the influence of the atmosphere and/or volume. Cefiderocol was evaluated only for the influence of the volume.

| Antibiotics | amikacin | gentamicin | ciprofloxacin | ceftazidime | meropenem | aztreonam | linezolid | cefiderocol |
| --- | --- | --- | --- | --- | --- | --- | --- | --- |
| *P. aeruginosa* | x | - | x | x | x | x | - | x |
| *K. pneumoniae* | x | - | x | - | x | - | - | x |
| *E. coli* | x | - | x | - | x | - | - | x |
| *A. baumannii* | x | - | x | - | x | - | - | - |
| *S. aureus* | - | x | x | - | x | - | x | - |

#### For the influence of the smaller final volume

Two types of plates were used: 96-well plates for tests with 100 and 200 µL final volume and 384-well plates for tests with 30, 50 and 100 µL in water-saturated atmosphere. For each condition, a minimum of three independent experiments were performed, each in triplicate. When available, MIC values were compared with the acceptable range of expected values for quality control strains, as defined in the EUCAST QC tables 14 (5) and AFST QC v7.0 (6). For bacteria, all tests (conditions are detailed in **Table 1**) were performed in Ca-MHB (except cefiderocol, for which MIC measurements were determined in ID-MHB) with an inoculum of 5.5 × 10^5^ CFU/mL and the plates were analyzed after 16-20 h of incubation at 37°C as recommended by EUCAST. For yeasts, the MIC for three antifungals (amphotericin B, fluconazole and micafungin) against both species were determined in supplemented RPMI 1640 with a final inoculum of 0.5-2.5 10^5^ CFU/mL. The plates were analyzed after 24 h of incubation at 37°C as recommended by EUCAST(4).

## Results

### Impact of the incubation atmosphere on evaporation of the medium

To compare evaporation, volumes were measured for all peripheral wells and for the wells in the middle of the plate. **Figure 1** shows that in an air atmosphere, 84 ± 2% of the volume remained in the corner wells, and that after 24 hours of incubation in 96-well plates, more than 90% of the volume remained in the edge wells. No volume loss was observed for the middle wells in the plate. For all wells, the final volumes remained stable in a water-saturated atmosphere (100% of the initial volume was found at 24 hours).

**Figure 1.**
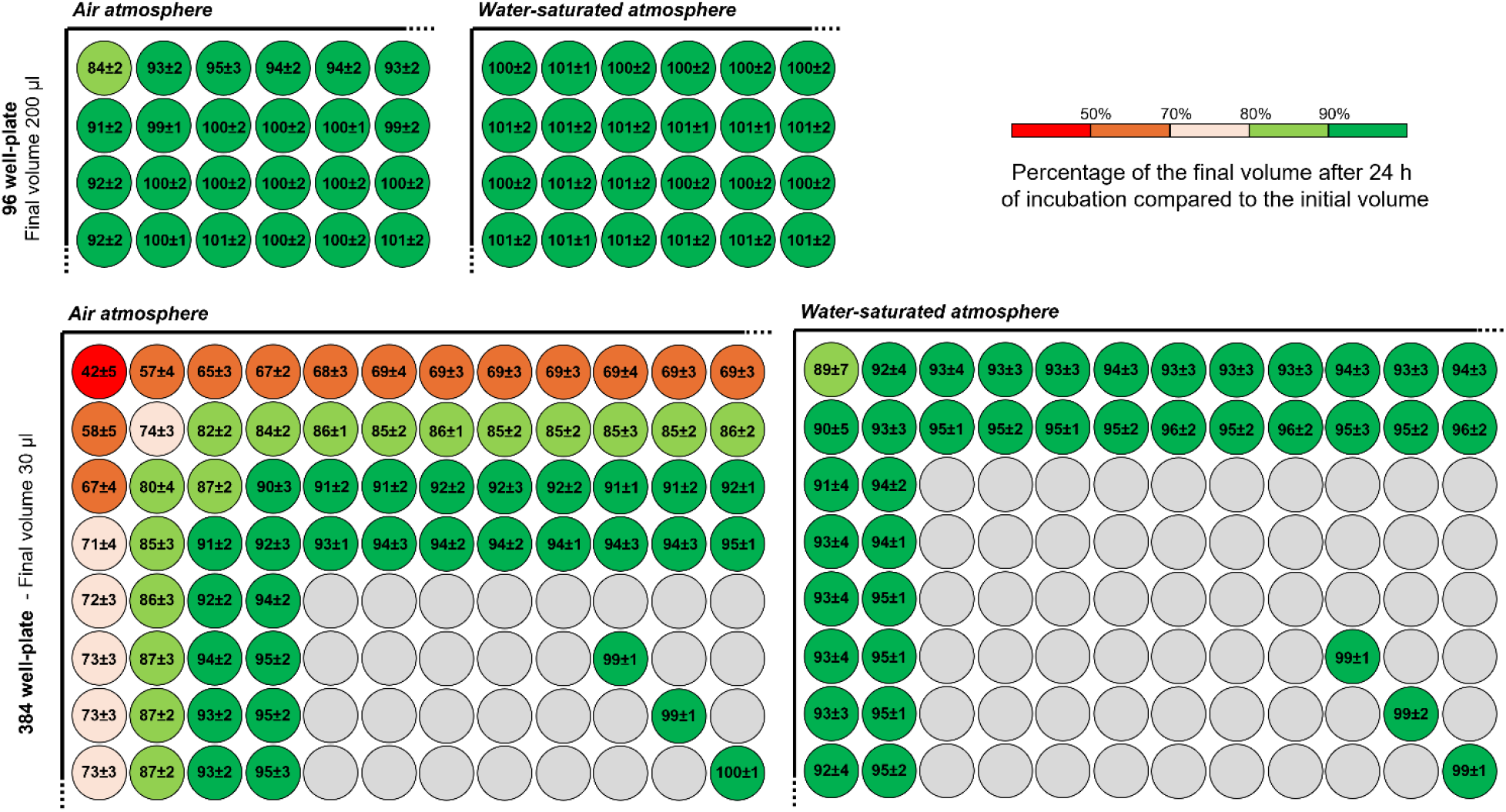
Percentage of evaporation at 24 h in air and water-saturated atmospheres. 96-well plate with final volume of 200 µL and 384-well plate with final volume of 30 µL were incubated for 24 h at 37°C without agitation. Results are expressed in percentage of final volume at 24 h compared to initial volume. The figure shows a schematic representation of the top left-hand corner of a microtiter plate.

For the 384-well plates with final volume of 30 µL, the effect was more pronounced, with only 42 ± 5% of the final volume remaining in the corner wells after 24 hours in an air atmosphere. For the edge wells (first column and first row), the final volume ranged from 57 ± 4% to 73 ± 3% at 24 hours, whereas the final volumes were close to 100% in the third column and third row, extending to the middle wells in the plate. For all wells in 384-well plates, the final volume remained above 90% in a water-saturated atmosphere. The results for all conditions tested (96-well versus 384-well plates and different volumes) are presented in Figure S1. The results were similar for the other areas of the corresponding plates. To limit the evaporation phenomenon, it was decided to maintain a water-saturated atmosphere for the remainder of the study.

### Impact of the incubation atmospheres on MIC values in 200 µL final volume

To ensure that a water-saturated atmosphere did not influence MIC measurement, the MIC values obtained in a final volume of 200 µL in both air and in a water-saturated atmosphere were compared. As illustrated in Figure 2, 90% (45 out of 50 measurements) of the MIC values fell within the acceptable ± 1 log2 dilution range, while only 5 values (10%) fell within the ± 2 log2 dilution range.

**Figure 2.**
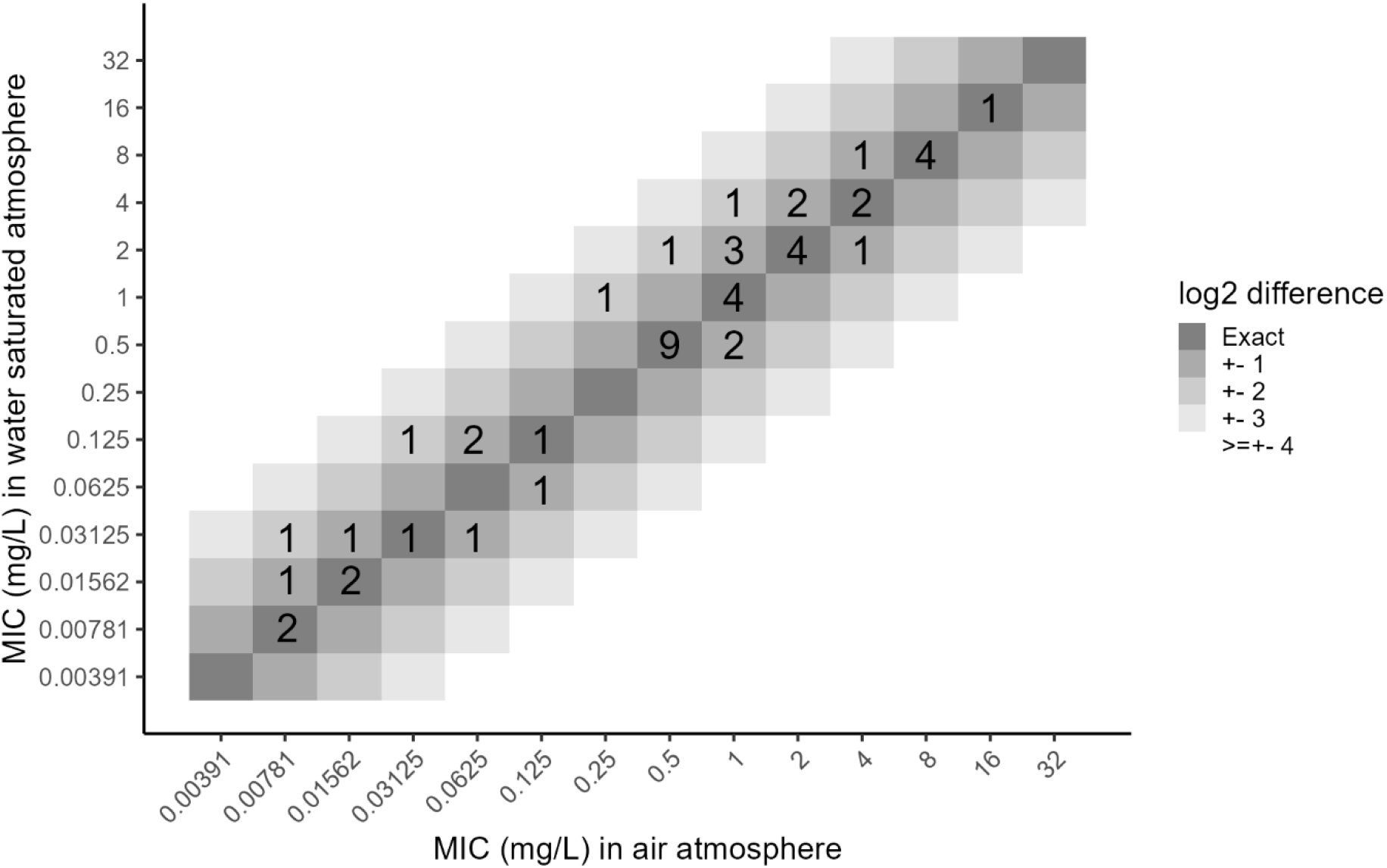
Comparison of MIC values obtained in 200 µL final volume in air or water-saturated atmosphere. MIC measurements were paired for both atmospheres and repeated in three independent experiments. The different grey levels shows differences of log 2 dilution from 0 (exact) to ≥ ± 4 log 2 dilutions.

### Impact of the volume reduction on MIC values

MIC measurements were then assessed in different volumes with different well plates for four gram-negative species (*Pseudomonas aeruginosa, Klebsiella pneumoniae, Escherichia coli and Acinetobacter baumannii*), one gram-positive (*Staphylococcus aureus*) and two Candida species (*Candida krusei, Candida parapsilosis*) for different antimicrobials representative of the different families used in therapy. Results are presented in **Figures 3, 4** and **S2**.

**Figure 3.**
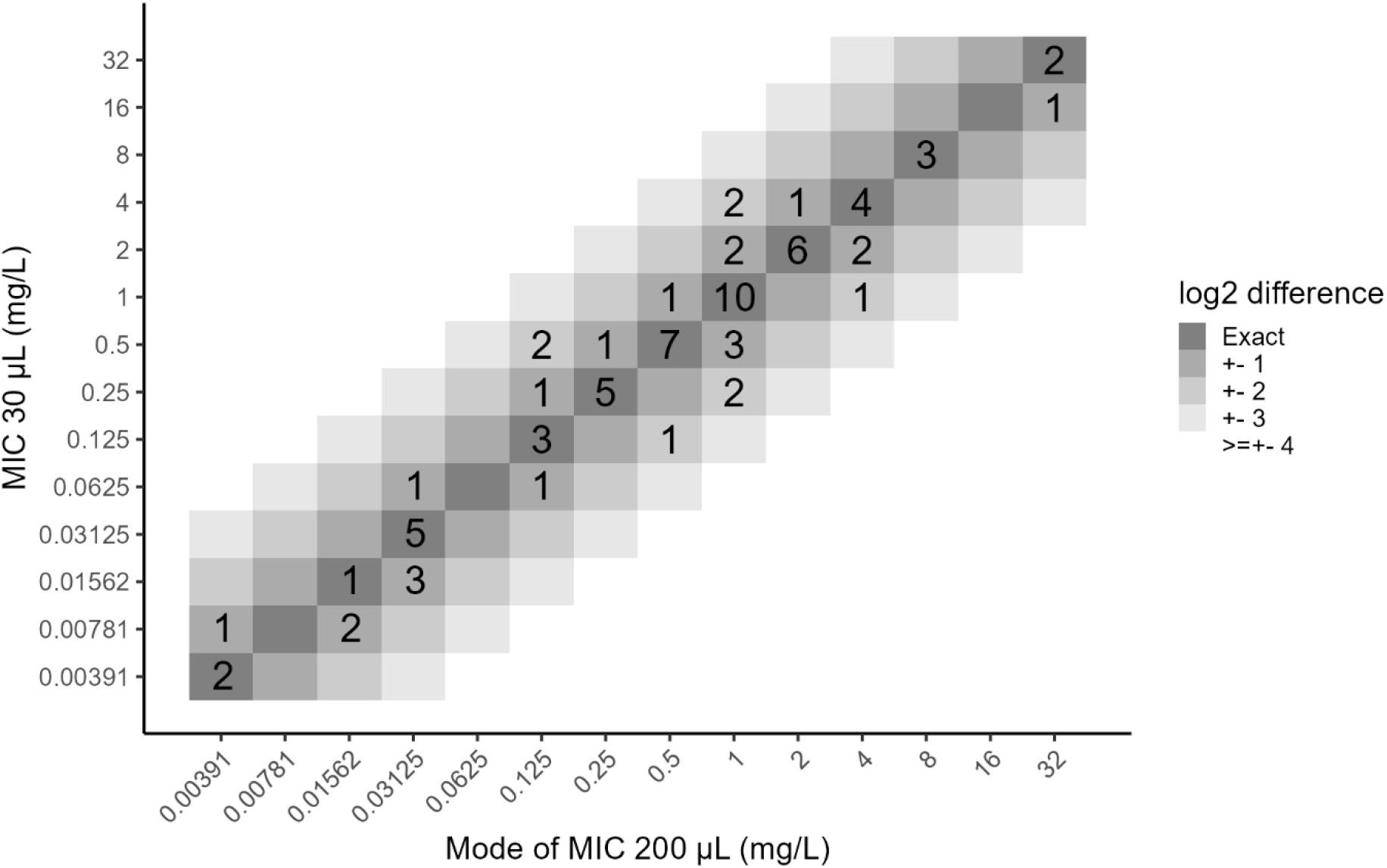
Comparison of MIC values obtained in 200 µL and in 30 µL final volume. MIC values of all antimicrobials against all bacterial and fungal species were plotted in differences of MIC values at 30 *µ*L against the mode of MIC values obtained at 200 *µ*L final volumes. The different grey levels shows differences of log 2 dilution from 0 (exact) to ≥ ± 4 log 2 dilutions.

**Figure 4.**
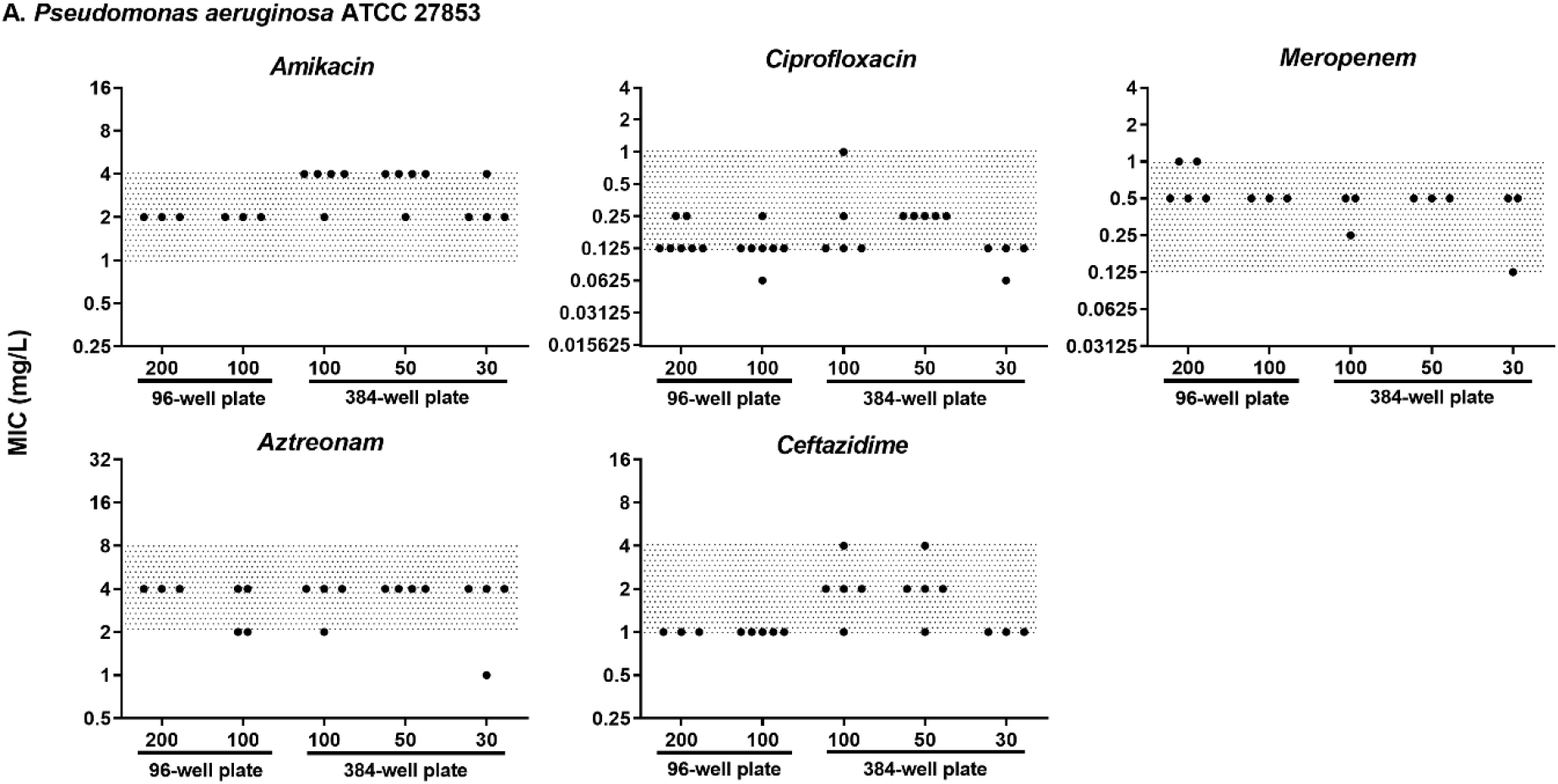
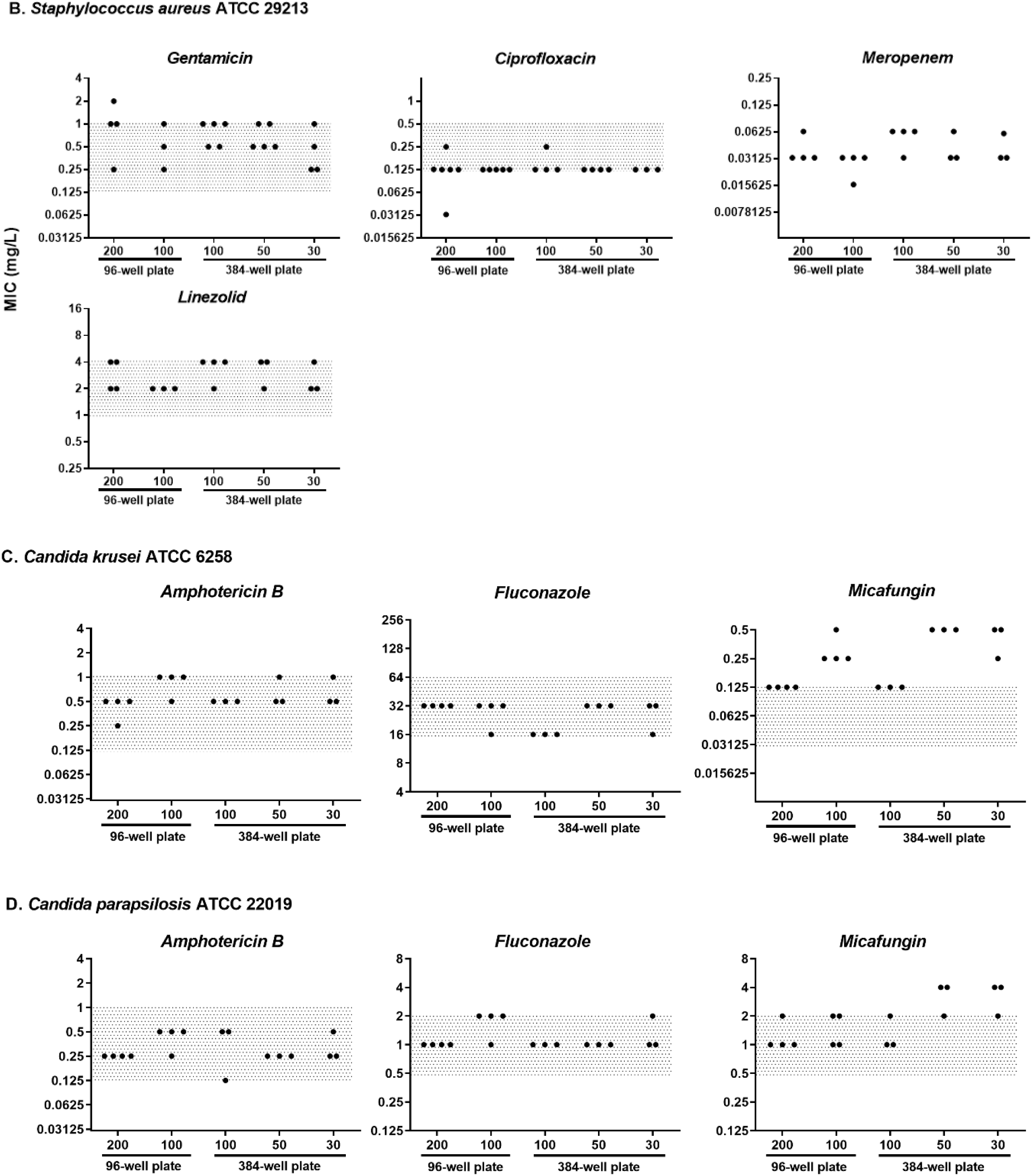
Comparison of MIC values depending on the different conditions. **A.** MIC values of amikacin, ciprofloxacin, meropenem, aztreonam and ceftazidime against *Pseudomonas aeruginosa* ATCC 27853. **B**. MIC values of ciprofloxacin, gentamicin, linezolid and meropenem against *Staphylococcus aureus* ATCC 39213. **C**. MIC values of amphotericin B, fluconazole and micafungin against *Candida krusei* and **D**. *Candida parapsilosis*. Each MIC was evaluated at least three independent times (black dot). The grey area represents the acceptable range for the respective quality control strains according to the EUCAST.

Figure 3 compares the MIC values obtained in 30 µL final volumes with the MIC values obtained in 200 µL, regardless of the species and antimicrobial tested. Most MIC values (48/76, 63.2%) showed an exact match between the MIC values obtained in both final volumes, while 20/76 (26.3%) showed a ±1 log2 dilution difference, representing 89.5% of the values that fell in the acceptable range of MIC. However, 8/76 (10.5%) values showed a ±2 log2 dilution difference. No MIC value differences greater than ±2 log2 dilutions were observed.

For *Pseudomonas aeruginosa* (**Figure 4A**), no significant difference was observed in MIC values, regardless of the volume used for the measurement. All MIC values were within the acceptable range (grey area) for the quality control strain, as defined by EUCAST. Comparable MIC values were obtained for *Staphylococcus aureus*, irrespective of the volume used for MIC evaluation (**Figure 4B**). For *Candida* sp. (**Figures 4C** and **4D**), results were highly comparable, with stable MIC values regardless of the final volume for amphotericin B and fluconazole. However, MIC values tended to be slightly increased by one to two dilutions for smaller volumes (30 and 50 µL) in the 384-well plates, with three out of three measurements for *C. krusei* and two out of three measurements for *C. parapsilosis* showing an increase for micafungin compared to the 200 µL volume in 96-well plates. The results for other gram-negative species (*Escherichia coli* ATCC25922, *Acinetobacter baumannii* ATCC19606, and *Klebsiella pneumoniae* ATCC43816) on a restricted selection of antibiotics are presented in **Figure S2** and showed no significant impact of the final volume on the MIC values of three different antibiotics: amikacin, ciprofloxacin, and meropenem. As reducing the volume for MIC measurement could have a major impact in terms of cost and the quantity of antimicrobial used, cefiderocol, for which expensive media (ID-MHB) is used, was also evaluated at 200 µL and 30 µL without showing any difference (**Figure S3**). On the other hand, a cost saving of around 87% was estimated when the volume was reduced for cefiderocol MIC evaluation (**Table S2**).

## Discussion

Optimizing MIC measurement conditions by reducing the final assay volume is a key challenge for future research. This approach aims to enable the testing of a larger number of strains while minimizing the costs associated with novel molecules, specialized media (which can be highly expensive), or synthesized compounds available only in limited quantities. In this study, we demonstrated that reducing the final assay volume to 30 µL generally does not significantly impact MIC values across various antimicrobials tested against Gram-negative and Gram-positive bacteria, as well as yeast species.

### The evaporation is more pronounced with small volume but can be counteracted in water-saturated atmosphere

A potential bias introduced by reducing the final assay volume is evaporation. In this study, we evaluated evaporation and found it to be more pronounced at smaller volumes (30 µL in 384-well plates), with the effect being most significant in corner and edge wells but rapidly becoming negligible in other wells. To mitigate evaporation in edge wells, we compared evaporation under identical conditions in microwell plates incubated in a water-saturated atmosphere, which significantly reduced the effect. Moreover, the MIC values of antibiotics measured in both air and water-saturated atmospheres were not significantly different (**Figure 2**). Therefore, to minimize evaporation at low final volumes, experiments should be conducted in a water-saturated atmosphere.

### Reducing the volume of experiments does not significantly influence MIC values

MIC measurements were then performed using standard volume in 96-well plates versus reduced volume in 384-well plates on a broad selection of bacterial and fungal species, as well as a wide range of antimicrobials. MIC values obtained under these different conditions were compared to the standard condition (96-well plates, 200 µL final volume) and were within the acceptable value range of the corresponding quality control strains as defined by EUCAST, when available (*P. aeruginosa, E. coli, S. aureus, C. parapsilosis* and *C. krusei*). Most MIC values fell within the acceptable ±1 log2 dilution range, indicating that the final volume and plate format did not significantly influence the measurement.

In our study, experiments with a final volume of 20 µL were also evaluated, but the MIC values were less reproducible (data not shown). While further volume reduction cannot be ruled out, the limit of reliable pipetting was reached with the pipettes used in this setting. Furthermore, although manual pipetting to fill the 384-well plates remains feasible, the use of a pipetting robot appears essential to improve reproducibility, as well as pipetting accuracy and speed.

The MICs for micafungin were at or above the acceptable limit. Micafungin is a sticky agent that tends to bind to plastics. A recent study (7) found that the number of tip changes during testing can influence the MIC values, probably due to the drug’s binding to plastic surfaces. This interaction could explain the discrepancies observed in our study. Since smaller volumes of micafungin were used in smaller wells, a greater proportion of the drug molecules may bind to the plastic, leading to a more significant impact on the MIC. This phenomenon is particularly pronounced when the MIC is low, as seen with a higher impact on *C. krusei* than on *C. parapsilosis*. These results suggest that for accurate comparisons of echinocandin MICs, laboratories should follow a consistent protocol, and that use of lower volumes may not be advisable.

### To reduce the volume of experiments, reduce materials needed and cost of the experiments

We then estimated the savings resulting from reducing the volume of experiments, from 200 µL in 96-well plates to 30 µL in 384-well plates.

Although the savings for a series of MIC measurements with ‘standard’ antibiotics (estimated for two strains against five antibiotics) are relatively modest, with a 15.1% cost reduction for 384-well plates with a 30 µL final volume (**Table S1**), they are considerably higher when expensive drugs requiring special media are used. In our example, for 10 bacterial strains with cefiderocol, cost savings could exceed 87% for 384-well plates with a 30 µL final volume (**Table S2**). Laboratory plastic consumption was estimated for an experiment on 100 strains, showing a usage of over 21.8 kg under standard conditions, versus 6.7 kg under reduced-volume conditions—corresponding to a 69.3% reduction in plastic waste (**Table S3**).

To our knowledge, this study is among the first to validate small-volume MIC testing based on the standards recommended by EUCAST or CLSI. While other studies have proposed miniaturizing MIC testing to increase experimental throughput, these approaches often rely on novel methodologies that require specialized (and potentially expensive) equipment. For instance, Needs *et al*. recently demonstrated good agreement with CLSI standards using a miniaturized broth microdilution antibiotic susceptibility testing (AST) method based on microcapillary devices for Gram-negative bacteria (8). Similarly, Choi *et al*. developed a rapid AST method using a single-cell approach combined with time-lapse microscopy imaging (9). However, in addition to their reliance on specialized equipment, these methods have limitations, particularly in their estimation of antimicrobial effects using a small number of bacteria. This may have overlooked the inoculum effect, which can significantly influence MIC values. (10). Previously, a study proposed an approach similar to ours for miniaturizing checkerboards, but tested only a single bacterial species on beta-lactam/beta-lactamase inhibitor combinations (11).

Overall, our study showed that reducing the final volume does not affect MIC values (except for micafungin) for a wide range of bacterial and yeast species across various antimicrobials with different mechanisms of action. On the other hand, reducing test volume can generate huge savings when assessing the activity of expensive molecules. Reducing the volume of MIC measurements would also make it possible to reduce the quantity of molecules needed, for example, in preliminary activity tests for new molecules that are difficult to synthesize or cannot be synthesized in large quantities.

## Acknowledgements

We would like to thank Agnès Audurier for her excellent technical assistance.

We also wish to thank Jeffrey Arsham, an American medical translator, for having reread and reviewed this original English-language article.

## Author contributions

J.C. and J.M.B. designed the project.

J.M., T.C. and E.B. conducted experiments.

J.C., J.M.B, K.B. and V.AC. conducted data analysis and drafted the manuscript.

J.C., K.B., S.M. and J.M.B. supported manuscript writing.

All authors reviewed the manuscript, performed final editing and approved the final version.

## Supplemental figures

Percentage of final volume after 24 h of incubation compared to the initial volume

**Figure S1.**
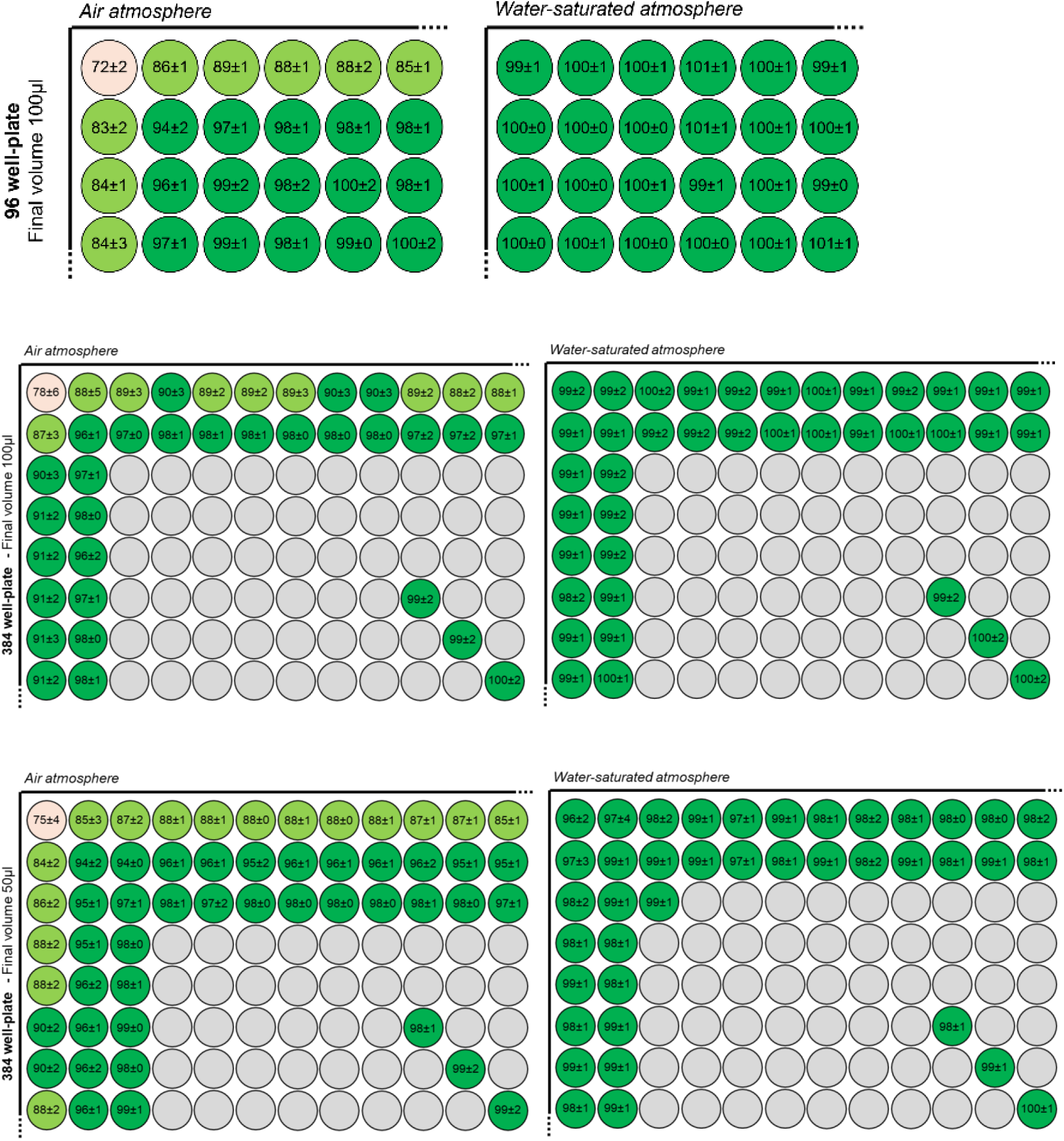
Percentage of evaporation at 24 h in air and water-saturated atmospheres. 96-well plate with final volume of 100 *µ*L and 384-plate with final volume of 50 and 100 *µ*L were incubated for 24 h at 37°C without agitation. Results are expressed in percentage of final volume at 24 h compared to initial volume. The figure shows a schematic representation of the top left-hand corner of a microtiter plate.

**Figure S2.**
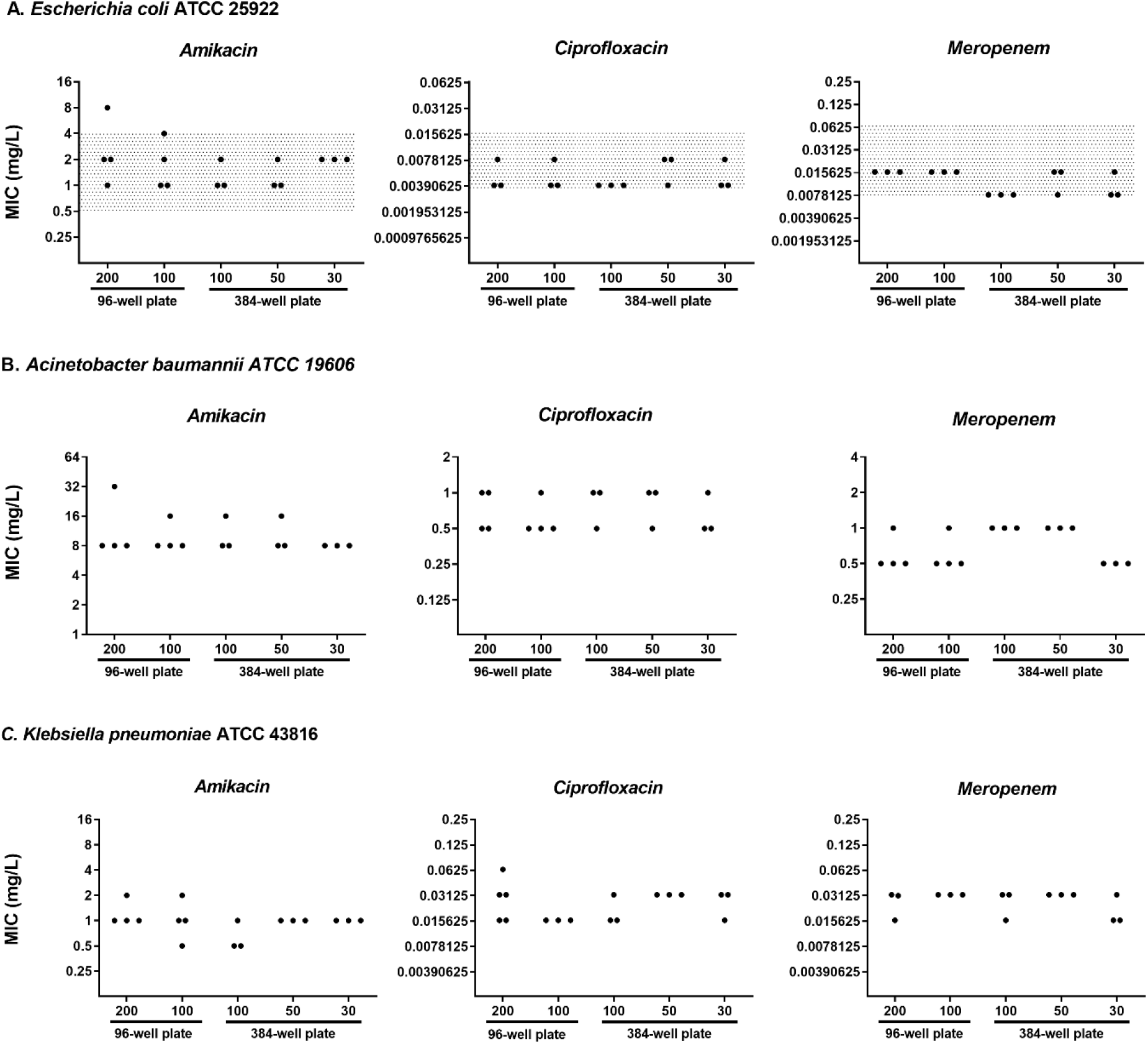
Comparison of MIC values for amikacin, ciprofloxacin and meropenem depending on the different conditions. **A**. MIC values against *Escherichia coli* ATCC 25922. **B**. MIC values against *Acinetobacter baumannii* ATCC 19606. **C**. MIC values against *Klebsiella pneumoniae* ATCC 43816. Each MIC has been evaluated at least three independent times (black dot). The grey area represents the acceptable range for the respective quality control strains according to the EUCAST when available.

**Figure S3.**
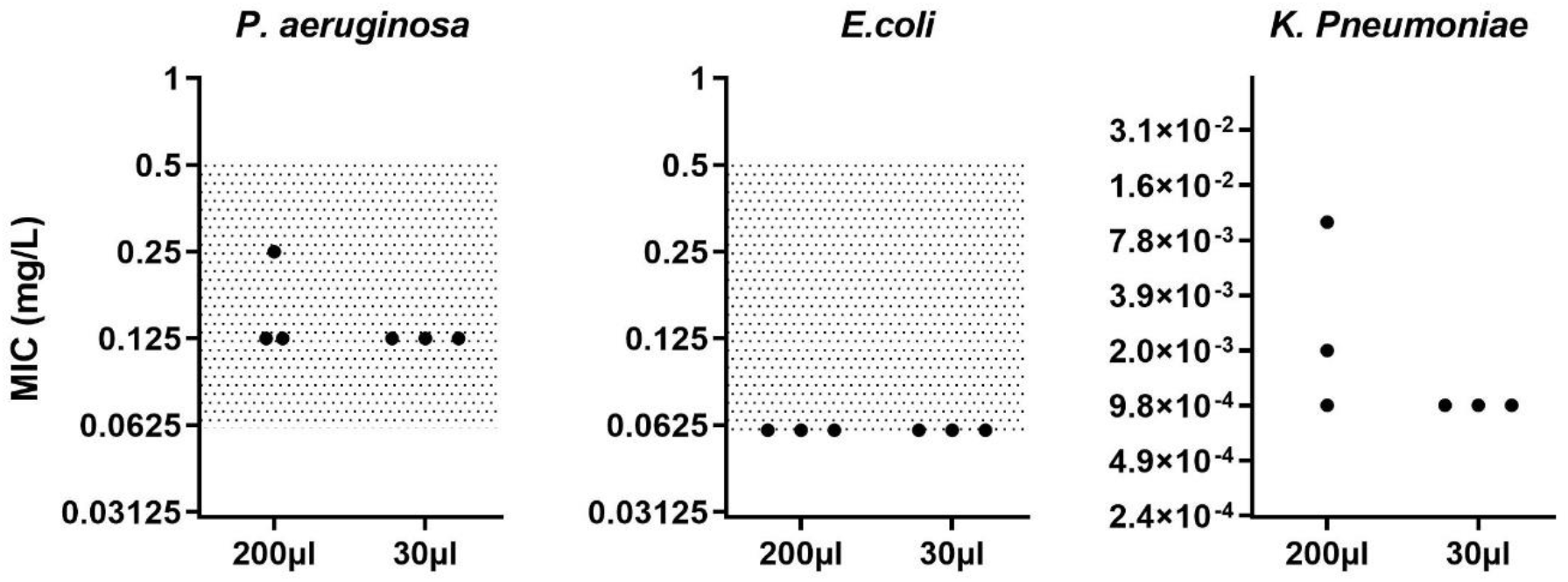
Comparison of MIC values for cefiderocol in water-saturated atmosphere depending on the different conditions against *Pseudomonas aeruginosa, Escherichia coli* and *Klebsiella pneumoniae*. Each MIC has been evaluated at least three independent times (black dot). The grey area represents the acceptable range for the respective quality control strains according to the EUCAST, when available.

**Table S1.** Estimation of the experimental cost for MIC measurements with 5 standard antibiotics against two bacterial strains.

| **Consumables**<br>(calculated for 2 strains with 5<br>antibiotics in triplicate) | **96 well-plate**<br>**Final volume 200 µL** | **384 well-plate**<br>**Final volume 30 µL** |
| --- | --- | --- |
| Plates | 5 | 1 |
| Dilution plate | 1 | 1 |
| Plates | 4 € (5 + 1 plates) | 8 € (1 + 1 plate) |
| Medium - MHB | 0,78 € | 0,20 € |
| Tips | 10,75 € | 14 € |
| Antibiotics : |  |  |
| ATM (590 €/g) | 3,55 € | 0,30 € |
| MEM (600 €/g) | 7,20 € | 0,60 € |
| AMK (18 €/g) | 0,21 € | 0,018 € |
| CIP (7,8 €/g) | 0,05 € | 0,004 € |
| CAZ (62 €/g) | 0,75 € | 0,06 € |
| **For one experiment** | **27.3 €** | **23.2 €** |
|  | 15.1% savings |  |

**Table S2.** Estimation of the experimental cost for MIC measurements with cefiderocol antibiotic against ten bacterial strains.

| **Consumables**<br>(calculated for 10 strains with 1<br>antibiotic in triplicate) | **96 well-plate**<br>**Final volume 200 µL** | **384 well-plate**<br>**Final volume 30 µL** |
| --- | --- | --- |
| Plates | 5 | 1 |
| Dilution plate | 1 | 1 |
| Plates | 4 € (5 + 1 plates) | 8 € (1 + 1 plate) |
| Medium – ID-MHB (210 €/L) | 42 € | 17 € |
| Tips | 10,75 € | 14 € |
| Antibiotics : |  |  |
| CFD (67000 €/g) | 3 350 € | 402 € |
| **For one experiment** | **3 406.75 €** | **441 €** |
|  | 87.1% savings |  |

**Table S3.** Estimation of Plasticware Consumption for MIC Measurements on 100 Strains.

| Consumables | 96 well-plate<br>Final volume 200 μL | 384 well-plate<br>Final volume 30 μL |
| --- | --- | --- |
| Plates | 15500 g (250 * 62 g) | 4000 g (50 * 80 g) |
| Dilution plate | 620 g (10 * 62 g) | 620 g (10 * 62 g) |
| Tips for antibiotic and inoculum preparation | 425 g (1700 * 0.25 g) | 425 g (1700 * 0.25 g) |
| Tips for dilution and spreading of bacterial inocula | 275 g (1100 * 0.25 g) | 275 g (1100 * 0.25 g) |
| Tips for plate preparation | 112 g (400 * 0.28 g)<br>4875 g (19500 * 0.25 g) | 1008 g (3600 * 0.28 g)<br>375 g (1500 * 0.25 g) |
| **Total** | 21807 g | 6703 g |

69.3 % savings
The weight of different plasticwares have been estimated as follows: 96-well plate = 62 g; 384-well plate = 80 g, 200 μL pipette tips = 0.25 g; 200 μL pipette tips for robot = 0.28 g

